# Osteopontin deficiency leads to the resolution of prostatic fibrosis and inflammation

**DOI:** 10.1101/2021.07.19.452973

**Authors:** Petra Popovics, Asha Jain, Kegan O. Skalitzky, Elise Schroeder, Hannah M. Ruetten, Mark Cadena, Kristen S. Uchtmann, Chad M. Vezina, William A. Ricke

**Affiliations:** Department of Urology, School of Medicine and Public Health, University of Wisconsin-Madison, Madison, WI; George M. O’Brien Center of Research Excellence, University of Wisconsin, School of Medicine and Public Health, Madison, WI; Department of Comparative Biosciences, School of Veterinary Medicine, University of Wisconsin-Madison, Madison, WI

## Abstract

Fibrogenic and inflammatory processes in the prostate are linked to the development of lower urinary tract symptoms (LUTS) in men. Our previous studies identified that osteopontin (OPN), a pro-fibrotic cytokine, is abundant in the prostate of men with LUTS and its secretion is stimulated by inflammatory cytokines potentially to drive fibrosis. This study investigates whether the lack of OPN ameliorates inflammation and fibrosis in the mouse prostate.

We instilled uropathogenic *E. coli* (UTI89) or saline (control) transurethrally to C57BL/6J (WT) or *Spp1*^*tm1Blh*^/J (OPN-KO) mice and collected the prostates one or 8 weeks later. We found that OPN mRNA and protein expression were significantly induced by *E. coli*-instillation in the dorsal prostate (DP) after one week in WT mice. Deficiency in OPN expression led to decreased inflammation and fibrosis and the prevention of urinary dysfunction after 8 weeks. RNAseq analysis identified that *E. coli*-instilled WT mice expressed increased levels of inflammatory and fibrotic marker RNAs compared to OPN-KO mice including *Col3a1, Dpt, Lum* and *Mmp3* which were confirmed by RNAscope.

Our results indicate that OPN is induced by inflammation and prolongs the inflammatory state; genetic blockade of OPN accelerates recovery after inflammation, including a resolution of prostate fibrosis.

## INTRODUCTION

Lower urinary tract symptoms (LUTS) developing secondary to prostate disease are a constellation of multiple urinary voiding and irritative symptoms that deteriorate the quality of life.^1^ The incidence of LUTS strongly correlates with age; approximately one in every four men aged 50 or above and more than 80% of men in their 80s have LUTS.^2^ Moreover, the current LUTS patient population is expected to grow due to the increasing life expectancy.^3^ Medical management of LUTS already poses an economic burden on our healthcare system, estimated to cost more than 4 billion dollars each year in the US.^4^ Medical treatments for LUTS target benign prostatic hyperplasia (BPH) and smooth muscle dysfunction, yet 45% of patients still progress to surgery due to treatment resistance and side effects^5,6^ which suggests the involvement of other pathological changes than what are currently recognized.

Prostates from LUTS patients exhibit markedly increased population of inflammatory cells, predominantly T lymphocytes and macrophages^7^, that led researchers to propose that BPH is an immune-mediated inflammatory disease.^8^ Inflammation has been shown to trigger proliferation and fibrosis, and has been associated with therapeutic failure.^9-11^ Histological inflammation also correlates with higher symptomatic index^12,13^, and patients with chronic inflammation have an increased risk of developing acute urinary retention.^14^ The etiology of inflammation is multifactorial, including autoimmune, hormonal, bacterial, viral and parasitic inducers.^9,15,16^ More than 50% of BPH prostates contain live bacteria with predominantly *Staphylococcus* (22%), *E. coli* (11%) and *Micrococcus spp* (8%).^17^ Several uropathogenic *Escherichia coli* (*E. coli*) strains have been isolated and transurethrally instilled in mice, resulting in marked inflammation, epithelial and stromal proliferation and fibrosis.^18-20^ This indicates that bacterial colonization may initiate the pathological cascade leading to BPH in at least a subset of LUTS patients.

Inflammation-induced fibrosis may be an important factor in symptom development but remains untargeted in the medical management of LUTS.^21^ LUTS are associated with increased stiffness in the periurethral prostate area^22^ and thicker collagen bundles compared to normal tissues, indicating fibrosis.^23^ Increased collagen accumulation also correlates with the clinical progression of LUTS.^24^ Treatments targeting androgen signaling and smooth muscle dysfunction may also perpetuate the fibrotic process, contributing to therapeutic failure.^25,26^ Consequently, our recent investigations focused on identifying targetable, inflammation-inducible factors of fibrotic signaling in the prostate.

Osteopontin (OPN) is a phosphoglycoprotein of the extracellular matrix with pro-inflammatory and pro-fibrotic properties.^27-29^ Our previous studies showed that there are significantly higher OPN protein levels in prostates from LUTS patients who progressed to surgery compared to incidental BPH tissue.^30^ We also identified increased mRNA expression of osteopontin (*Spp1* gene) during a qPCR screen in carrageenan-induced prostatic inflammation in rats.^31^ OPN is secreted by immune cells and prostate epithelial and stromal cells and stimulated by the inflammatory environment.^30,32^ Using prostatic stromal cells, we demonstrated that OPN promotes the expression of cytokines such as *IL6, CXCL1*-*2* and *8*, leading us to propose an intraprostatic exacerbation loop of inflammation driven by the increasing levels of OPN.^30^ This suggests that upregulation of OPN in prostatic inflammation may drive the transition to a chronic condition and, consequently, augment fibrosis.

The present study aims to test the requirement for OPN in inflammation-mediated fibrosis. Using an established *E. coli*-induced prostatic inflammation model^18^, we demonstrated that OPN mRNA and protein levels increase in response to inflammation. Using OPN knockout mice (OPN-KO), we showed that inflammation and fibrosis resolve in the absence of OPN, but OPN is not required for the development of acute inflammation. Bacterial instillation also increased voiding frequency in wild type mice but not in OPN-KO mice. Resolution of fibrosis in OPN-KO mice was consistent with the downregulation of pro-inflammatory and pro-fibrotic genes, such as *Lum, Dpt, Col3a1* and *Mmp3*. These results indicate that OPN may elongate the healing process after an inflammatory attack in the prostate and contribute to the development of fibrosis-related urinary symptoms.

## RESULTS

### Prostatic OPN expression is induced in *E. coli*-instilled mice

Our first goal was to establish the localization pattern of OPN in the mouse prostate and to determine whether OPN expression is induced by inflammation. We selected a transurethral instillation method using the UTI89 *E. coli* strain that was previously determined to trigger robust prostatic inflammation and fibrosis one week after the first of two transurethral instillations.^18^ The study establishing the model determined that instillation of 100 µl fluid, the same volume used in our experiments, specifically distributed to the dorsolateral prostate and *E. coli* at OD 0.8 colonized the dorsolateral and the anterior prostate. Our preliminary tests identified signs of inflammation mainly in the dorsal (DP) and the ventral prostate (VP) lobes; thus, we selected these lobes for further investigation. Using the RNAscope *in situ* hybridization technology, we identified that the *Spp1* gene, which encodes OPN, has a low expression level in saline-instilled control mouse prostates and is primarily localized to epithelial cells (Fig. 1A and 1C., Fig. S1). However, in the inflamed prostate, *Spp1* expression undergoes a robust increase with dense intracellular pattern in cells. This upregulation is likely due to the high expression of *Spp1* in various immune cells.^33,34^ In many cases, increased *Spp1* expression was associated with the expansion of prostate glands by several layers of proliferating epithelial and infiltrating inflammatory cells (Fig. S1). We found that the *Spp1* particle number was 30-fold elevated in the DP (p<0.01) but not significantly different in the VP. We also investigated the protein levels of OPN in these tissues using immunohistochemistry (IHC) and found a 1.77-fold elevation (p<0.001) in the DP but no significant change in the VP (Fig. 1B, 1D, Fig. S2). This corresponds to earlier reports^18^ and our subsequent observation, that there is significant inflammation in the DP one week after bacterial instillation but not in the VP.

**Figure 1:**
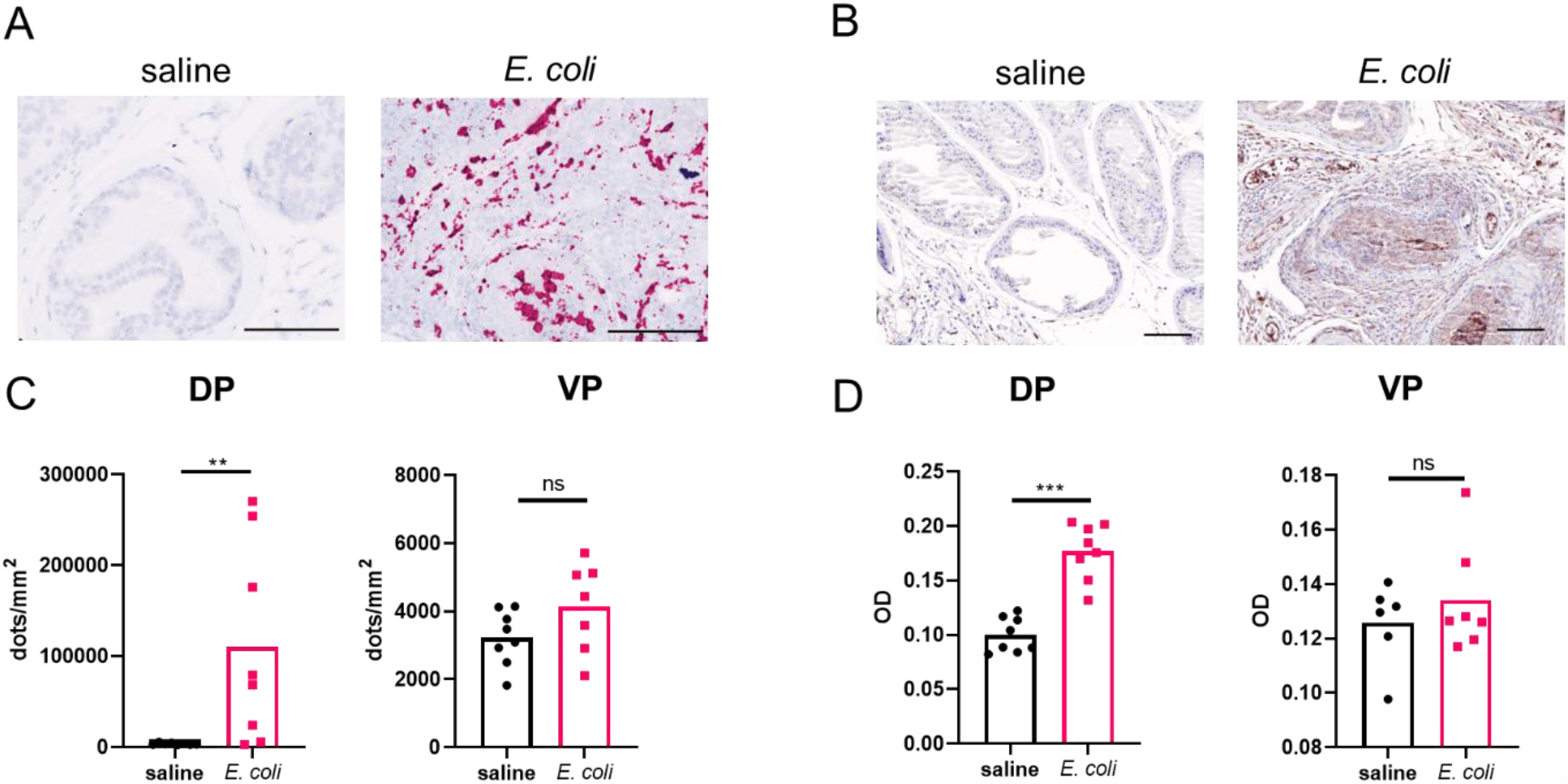
SPP1 gene expression and OPN protein levels are induced in bacterial prostatic inflammation. Both mRNA and protein expression significantly increases in response to bacterial instillation in the dorsal (DP) but not in the ventral prostate (VP). Panel **A** and **C** show RNAscope staining and signal quantification, respectively. Images were captured at 40x magnification. Panel **B** and **D** show immunohistochemistry staining and optical density, respectively (OD). RNAscope signal was normalized to tissue area and OPN OD was quantified by inForm software. Images were captured at 20x magnification. Scale represents 100 µm. ^**^:p<0.01

### Urinary function is improved in OPN-KO mice

To identify the role of OPN in bacteria-induced urinary dysfunction and prostatic inflammation, we utilized an OPN knockout strain, in which exons 4-7 were disrupted in all cells by the insertion of a neomycin cassette^35^, and compared to wild-type (WT) mice instilled simultaneously. In addition to examining the model in the acute inflammation phase at week one, we also investigated the chronic inflammatory state 8 weeks after bacterial instillation (Fig. 2A). Controls from both mouse strains were instilled with saline. To determine whether WT and OPN-KO strains have differences in colonization rates, we collected free-catch urine from *E. coli*-instilled mice 24 hours after the second instillation and prepared serial dilutions for subsequent plating on kanamycin agar plates. From both WT and OPN-KO strains, we identified one case from each mouse strain to have not been colonized by *E. coli* in our cohort. For the rest of the cases, the number of colony-forming units (CFUs, Figure 2B) were not statistically different between the two mouse strains, providing evidence that histological and functional changes investigated in the next steps are not related to altered colonization rates. The high variability in CFU counts (zero or 10^5^-10^9^) also highlights that the model may produce great variations of inflammatory levels in the prostate in both strains.

**Figure 2:**
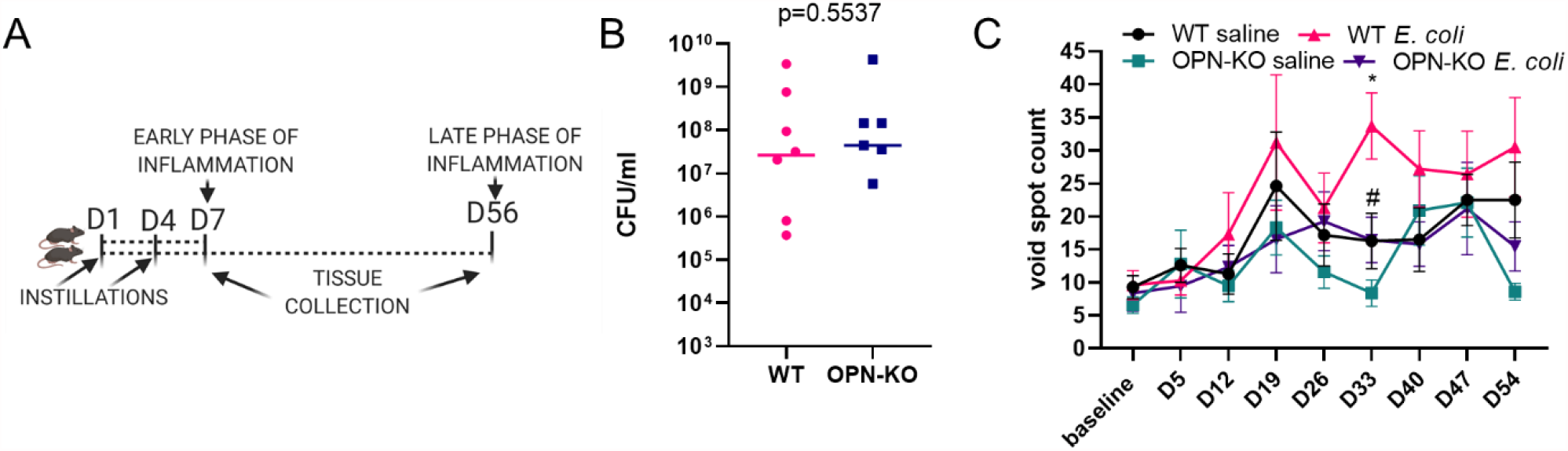
*E. coli*-induced urinary dysfunction is ameliorated in OPN-KO mice. Panel **A** shows experimental set up for the acute and chronic inflammatory time points. WT or OPN-KO mice were transurethrally instilled with *E. coli* or saline two times, 3 days apart and were euthanized at day 7 (acute inflammation) or 56 (chronic inflammation). Panel **B** shows that colony forming units (CFUs) in free-catch urine were not significantly different between WT and OPN-KO mice 24 hours after instillation. One data point from each group were zero and were not added to the figure but were included in the statistical analysis. Panel **C** shows void spot counts sized 0-0.1 cm^2^ measured weekly throughout the course of the chronic experimental set up. Significant differences were found WT saline vs. WT *E. coli* (^*^) and WT *E. coli* vs. OPN-KO *E coli* (#) at day 33 using Mann-Whitney test. ^*^,#: p<0.05.

Our next goal was to test whether OPN mediates inflammation-induced urinary dysfunction. *E. coli* can colonize and cause inflammation in multiple organs of the urinary tract including the bladder, prostate lobes, kidneys^18,36^ and the prostatic urethra (unpublished observation). Bacteria-induced urinary dysfunction may develop independently from prostatic inflammation, since it also develops in female mice^36^. Thus, in this model, urinary functional changes are the consequences of multiple pathological disturbances in the urogenital system. Earlier studies have established that transurethral instillation of gram-positive bacteria strains and *E. coli* induce urinary dysfunction in the form of increased urinary frequency.^37,38^ To determine whether void spot numbers increase, as an indicator of urinary dysfunction, over the course of the 8-week recovery period after *E. coli* instillation, we performed weekly void spot assays and analyzed the number of urine spots in the 0-0.1 cm^2^ category. Interestingly, inflammation increased void spots at day 33 in WT but not in OPN-KO mice relative to saline controls (Fig. 2C). Comparing only the two *E. coli*-instilled groups identified a significantly lower void spot count at day 33 in OPN-KO mice (Fig. 2C). It was previously shown that the UTI89 strain increases urinary output, measured as an increase in the total area of urine.^18^ However, this did not occur in our experiment (data not shown) possibly due to the difference in VSA timepoint (day 5 instead of day 7) or our slight modification of the bacterial propagation method, which involved shaking instead of the static culture used previously.^18^ In conclusion, in this study, WT mice developed a temporary dysfunction after 33 days which was prevented in OPN-KO mice.

### Bacteria-induced inflammation is ameliorated in OPN-KO mice

Histological inflammation was evident in the DP of WT and OPN-KO mice at day 7 post inflammation, whereas the histology of the VP remained visually unaffected (Fig. 3). In contrast, a high degree of inflammation was sustained in the DP and also spread to the VP in WT mice, while prostatic histology was more similar to the saline control in OPN-KO at this time point. Next, we quantified inflammatory levels by labeling CD45+ cells in the different experimental groups (Fig. 4). One week after the instillation, there was a significant increase in the number of CD45+ cells in the DP in both WT and OPN-KO mice compared to their respective saline controls (Fig. 4A and 4C). In the VP, CD45+ count was only significantly elevated in the OPN-KO mice (Fig. 4D). There was no significant difference between WT and OPN-KO *E. coli*-instilled mice in the VP (Fig. 4D). At day 56 however, there was a significant increase in CD45+ cells only in WT mice in the DP, but not in OPN-KO mice, compared to saline controls (Fig. 4B and 4E). There was also a significant reduction in OPN-KO *E. coli*-instilled mice when directly compared to WT (Fig. 4E). In the VP, CD45+ cell densities were not significantly different in WT E. coli vs. WT saline at day 56 using Kruskal-Wallis test (Fig 4F), but were significant using direct comparison with Mann-Whitney test (data not shown, p=0.0350). There was no significant change in OPN-KO VP CD45+ cells compared to saline controls or WT *E. coli* (Fig. 4F). CD45+ cell counts were variable within experimental groups similarly to infection rates shown earlier in this study.

**Figure 3:**
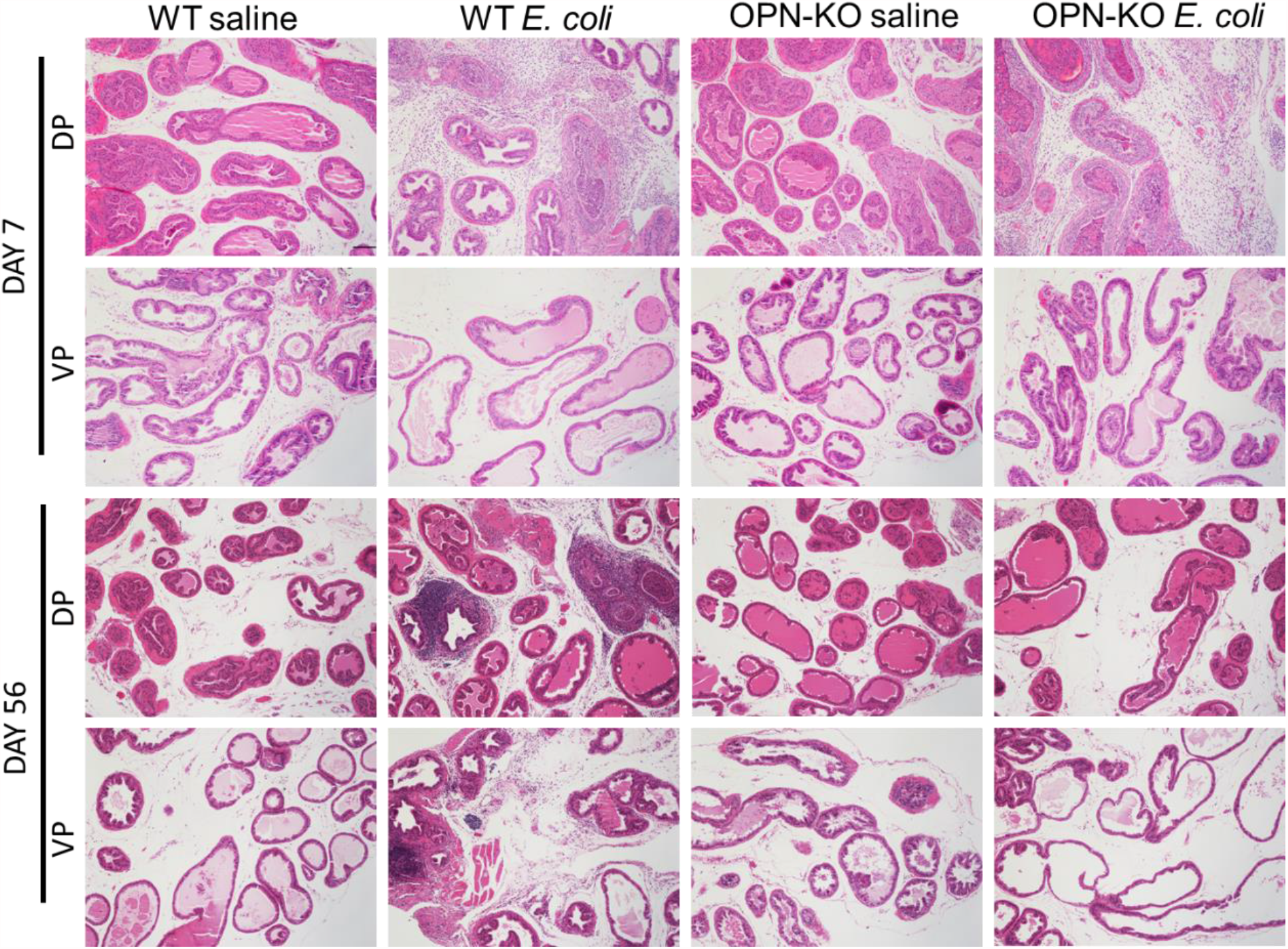
Histological inflammation resolves in OPN-KO mice. Seven days after the first E. coli instillation the stroma is occupied by inflammatory cells and multiple layers of epithelial and smooth muscle cells develop in both WT and OPN-KO mice. No visible inflammation is seen in the VP. After 2 months, inflammatory cells persist in WT mice in the DP and also infiltrate the VP, but the normal prostate histology is restored in the majority of OPN-KO mice. Images were captured at 20x magnification. Scale represents 100 µm.

**Figure 4:**
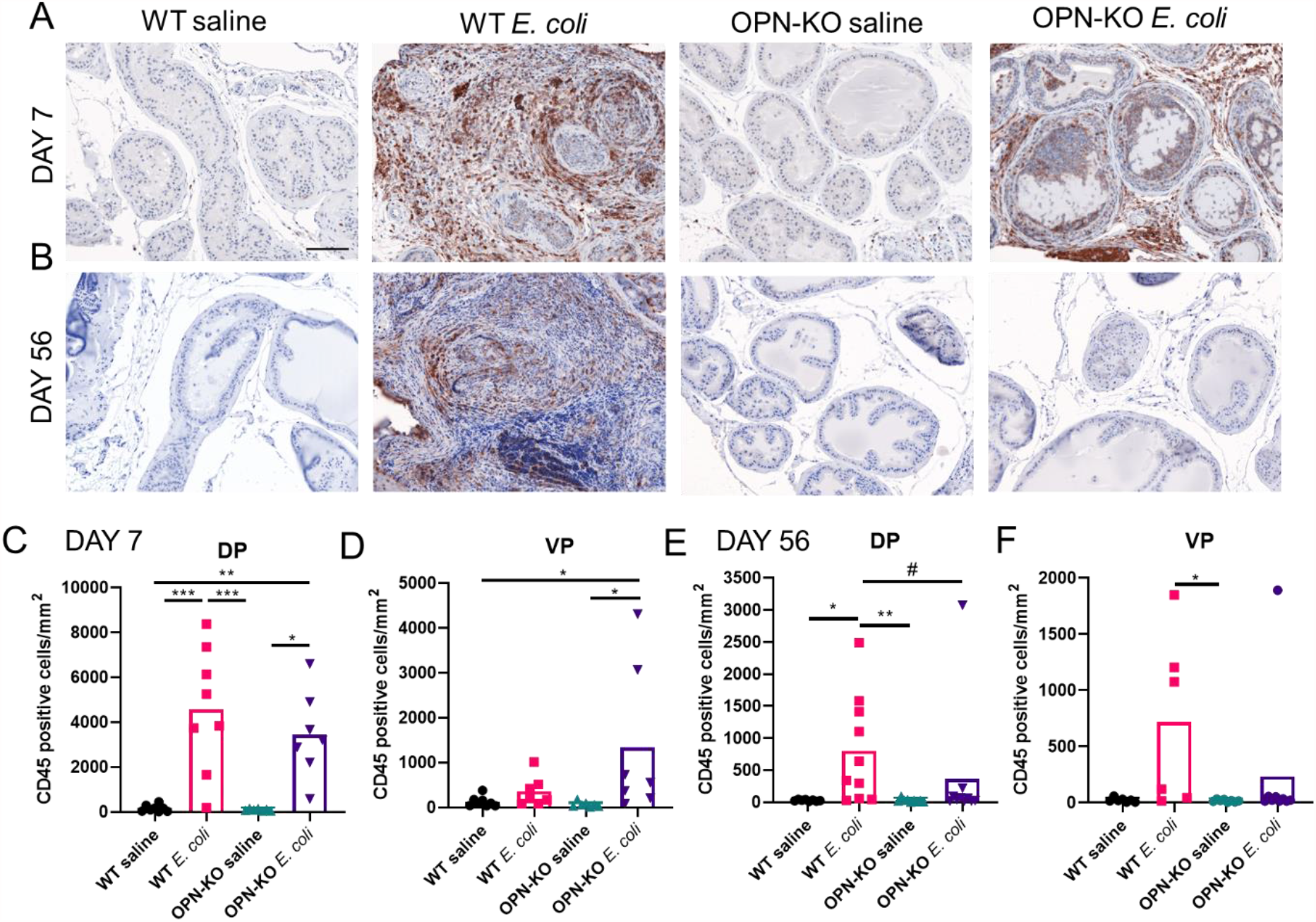
Inflammatory cell numbers decrease over time in OPN-KO mice. CD45+ cells were identified using immunohistochemistry and were counted and normalized to tissue area. Panels **A** and **B** show representative images of CD45 staining 7 and 56 days after the first *E. coli*- or saline instillation. Panels **C** and **D** show comparison of CD45+ counts across experimental groups in the DP and the VP at day 7. Panels **E** and **F** show comparison of CD45+ cell numbers in saline and *E. coli*-instilled WT and OPN-KO mice 56 days after the initiation of the experiment. Images were captured at 20x magnification. Scale represents 100 µm. ^*^: multiple analysis with Kruskal-Wallis test, #: pair-wise comparison with Mann-Whitney test between WT *E. coli* and OPN-KO *E. coli* mice. ^*^,#: p<0.05; ^**^: p<0.01; ^***^: p<0.001.

### Chronic accumulation of collagen is suppressed in OPN-KO mice

To explore potential alterations in inflammation-related prostatic fibrosis in OPN-KO mice, we examined collagen abundance by using PSR staining and imaging through polarized light filters.^23^ There was no significant change in collagen abundance one week following bacterial instillation in either WT or OPN-KO mouse strains (Fig. 5A, 5C and 5D). Colors observed through polarization imaging of collagen birefringence can imply a change in fiber size, alignment and packing, cross-linking, interstitial ground substance, and water content.^39^ We identified a significant decrease in the proportion of green fibers and an increase in yellow fibers in WT mice DP (Fig. S3), indicating a higher degree of collagen remodeling in this group in response to *E. coli* instillation. Two months after instillation, there was a 3-fold and a 3.3-fold increase in total collagen content in the DP and the VP in *E. coli*-instilled WT mice, respectively, and was significantly lower in OPN-KO mice vs. WT mice (Fig. 5B, 5E and 5F). We found no significant change in the distribution of colors in the chronic inflammation phase (Fig. S3).

**Figure 5:**
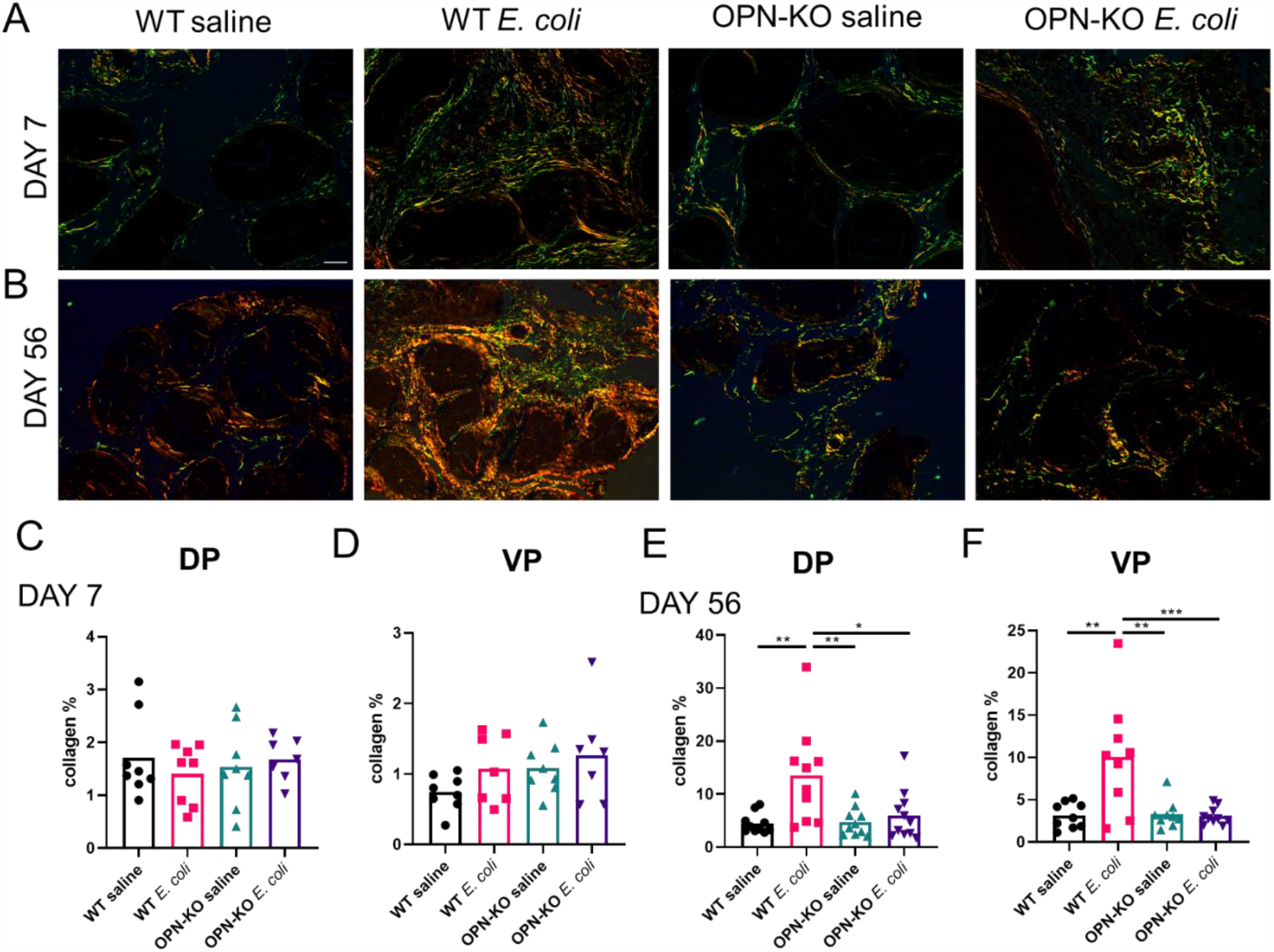
The inflammation-induced surge in collagen abundance is suppressed in OPN-KO mice. Panel **A** shows that signs of an increase in collagen abundance were observed in the DP one week after *E. coli*-instillation, but quantification of collagen % did not identify significant changes in either the DP or the VP (**C** and **D**). Panel **B** shows an example of the DP two months after infection, and Panels **E** and **F** identify significant increase in collagen content and reduction in OPN-KO mice in both the DP and the VP. Significance was calculated using One-way ANOVA with Tukey’s post-hoc test. Images were captured at 40x magnification. Scale represents 100 µm. ^*^: p<0.05; ^**^: p<0.01; ^***^: p<0.001.

To assess changes specifically in the abundance of collagen-I, we detected the protein expression of alpha-1 type I collagen subunit (Col1a1) using IHC and evaluated its optical density.

This method identified that inflammation significantly increases prostatic collagen-I level in both WT and OPN-KO mice in the acute model (Fig. 6A, 6C and 6D). On the other hand, Col1a1 abundance was only increased in WT DP and VP in infected mice compared to saline controls two months after bacterial instillation, and was significantly reduced in OPN-KO *E. coli* vs. WT *E. coli* prostates (Fig. 6B, 6E and 6F).

**Figure 6:**
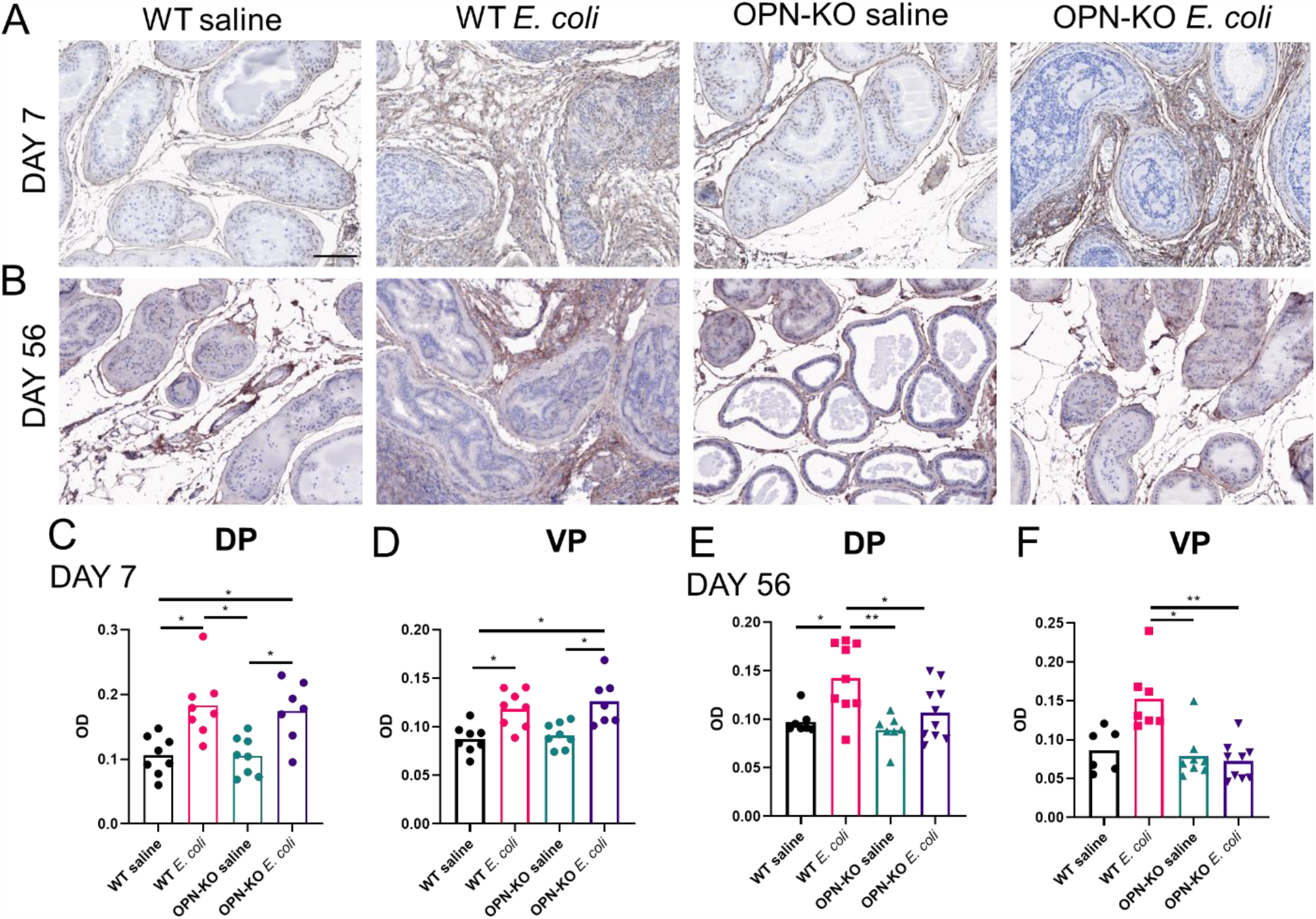
Inflammation-induced Col1a1 accumulation is ameliorated in OPN-KO mice. Panel **A** shows representative examples and **C** and **D** demonstrate that Col1a1 optical density (OD) increases in both WT and OPN-KO mice one week after bacterial instillation. Panel **B** demonstrates that Col1a1 abundance is visibly increased in WT mice, but not in OPN-KO mice 2 months after *E. coli*-instillation. Quantification of Col1a1 OD in the DP and the VP is shown in panels **E** and **F**, respectively. Significance was tested using Kruskal-Wallis test. Images were captured at 20x magnification. Scale represents 100 µm.

### Identification of pro-inflammatory and pro-fibrotic genes contributing to OPN action

To identify differentially expressed genes (DEGs) associated with bacterial inflammation and the loss of OPN expression, we performed bulk RNA-seq of the VP from 3 mice/group. Tissues with the highest collagen abundance were selected from *E. coli*-instilled WT and OPN-KO mice. We used paired-end 150b RNA-seq with >30 million read sequencing depth. The data was trimmed, quality checked, and mapped to the mouse genome. The unsupervised multidimensional scaling plot comparing all 4 experimental groups highlighted that 2 of the 3 samples from the WT *E. coli*-instilled group clustered differently from the rest of the samples (Fig. 7A). The transcriptomic variation within the WT *E. coli* group is likely due to the high degree of histological diversity seen in this model which can alter cell type dominance, but this also indicates that the most robust differences will be identified in this group, as expected. The bioinformatics analysis was focused on two direct comparisons: WT saline vs. WT *E. coli* and WT *E. coli* vs. OPN-KO *E. coli* to determine the effects of 1.) bacterial instillation and 2.) the loss of OPN expression. Hierarchical clustering analysis also confirmed that 2 out of 3 samples from the WT *E. coli* group differ in transcriptomic signature from the WT saline and OPN-KO *E. coli* samples (Fig. 7B and 7C). Since immune cell markers were over-represented in top DEGs due to high inflammatory cell infiltration, we focused on selecting a list of pro-inflammatory and pro-fibrotic genes with significant expressional changes and potential implication in maintaining the chronic fibrotic process (Fig. 7D, Table S1/<u>will be available after publication</u>). These DEGs in the WT *E. coli* vs. WT saline included genes that encode collagens (*Col1a1, Col3a1, Col5a1, Col5a2, Col5a3, Col6a5, Col14a1* and *Col15a1*), collagen processing enzymes (*Lox, Loxl1, Loxl2* and *Loxl3*), non-collagenous extracellular matrix (ECM) structural proteins (*Eln, Dpt, Dcn, Fbn1* and *Matn2*), proteoglycans (*Lum*), cell surface glycoproteins (*Dsc2*), proteases (*Habp2, Mmp3*, -*7*, -*9*, -*12* and -*23*), growth factors and binding proteins (Igf1 and Igfbp4), and cytokines and chemokines (*Tnf, Il1b, Il16, Il33, Cxcl1*, -*2*, -*12*, -*13*, -*17, Ccl2*, -*8*, -*9* and -*20*). Vimentin, a marker of stromal ^40^ and inflammatory cells^41^ was also increased, indicating the expansion of the stromal compartment and inflammation. We also observed a decrease in a few DEGs, most importantly in Cyp2b10b, which encodes a cytochrome P450 enzyme involved in xenobiotic and steroid hormone metabolism. Interestingly, none of the 46 genes was upregulated in *E. coli*-instilled OPN-KO mice vs. OPN-KO saline controls (Fig S4, Table S2/<u>will be available after publication</u>), and Habp2 was even suppressed in this comparison.

**Figure 7:**
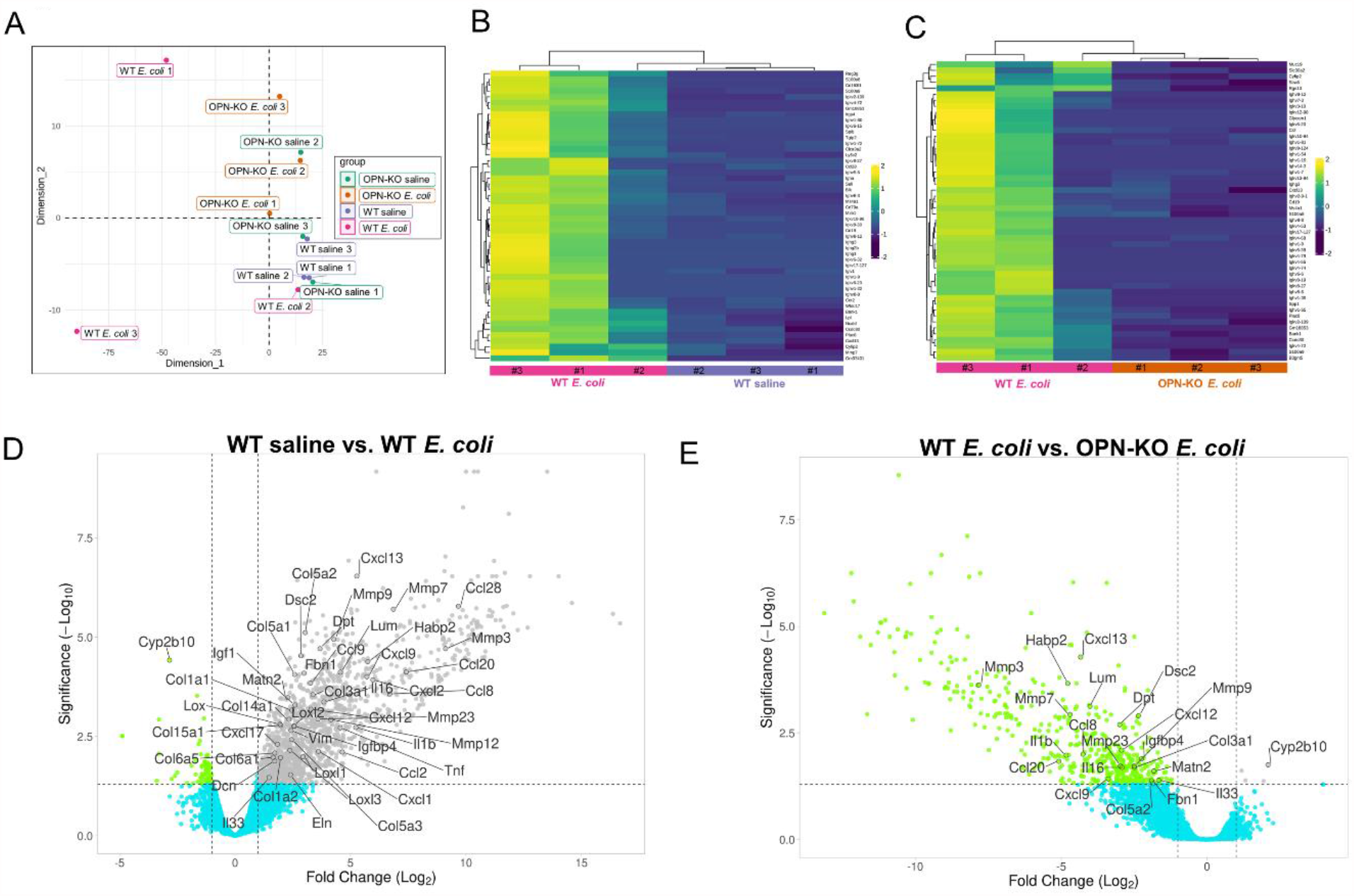
OPN deficiency obliterates the chronic inflammatory and pro-fibrotic gene expression signature related to *E. coli* infection. Panel **A** shows an unsupervised multidimensional scaling plot based on RNA-seq of triplicate samples from WT saline, WT *E. coli*, OPN-KO saline and OPN-KO *E*.*coli* experimental groups. This plot demonstrates that two of the three WT *E. coli* samples differ greatly from the rest of the tissues. Similarly, hierarchical clustering analysis shown in Panels **B** and **C** with WT saline vs. WT *E. coli* and WT *E. coli* vs. OPN-KO *E. coli* contrasts demonstrate that 2 tissues of the WT *E. coli* group cluster differently, which is likely due to a more robust inflammation developed in these samples compared to the third tissue. Panel **D** shows a volcano plot with selected pro-inflammatory and pro-fibrotic DEGs that are significantly increased in response to bacterial instillation in WT mice. We also highlighted a gene, Cyp2b10, that is downregulated in this condition. Panel **E** identifies which of the selected genes are downregulated (or upregulated in the case of Cyp2b10) in OPN-KO *E. coli* mice.

The upregulated 46 genes were then assessed for significant changes in the OPN-KO *E. coli* vs. WT *E. coli* contrast, providing a list of 21 DEGs that were downregulated in OPN-KO mice (Fig. 7E, Table S3<u>/will be available after publication</u>). These included collagens (*Col3a1* and *Col5a2*), other ECM genes (*Lum, Dpt, Fbn1* and *Matn2*), proteases (*Mmp3*, -*7*, -*9*, -*23* and *Habp2*), cytokines and chemokines (*Il1b, Il16, Il33, Cxcl9, Cxcl12, Cxcl13, Ccl8* and *Ccl20*), and others (*Dsc2* and *Igfbp4*). This indicates that *E. coli*-induced pro-inflammatory and pro-fibrotic processes were suppressed or ameliorated in mice lacking OPN.

### Validation of selected fibrosis markers

To test the validity of RNAseq results, we selected 4 genes (*Col3a1, Lum, Dpt* and *Mmp3*) to assess expressional differences in both the VP and the DP between WT *E. coli* and OPN-KO *E. coli* groups (Fig. 8). Expressional data for all four experimental groups and representative images are included in the supplemental data (Fig. S5). We found that all four genes were exclusively expressed in the stromal compartment. *Col3a1* expression was significantly downregulated in the VP and downregulation in the DP of OPN-KO *E. coli* mice approached significance (p=0.0653). *Lum* and *Dpt* expression was also significantly lower in OPN-KO *E. coli* mice. Interestingly, *Mmp3* expression was very limited and only occurred in prostates with very high inflammatory rate mainly in the WT *E. coli* group (Fig. S5)

**Figure 8:**
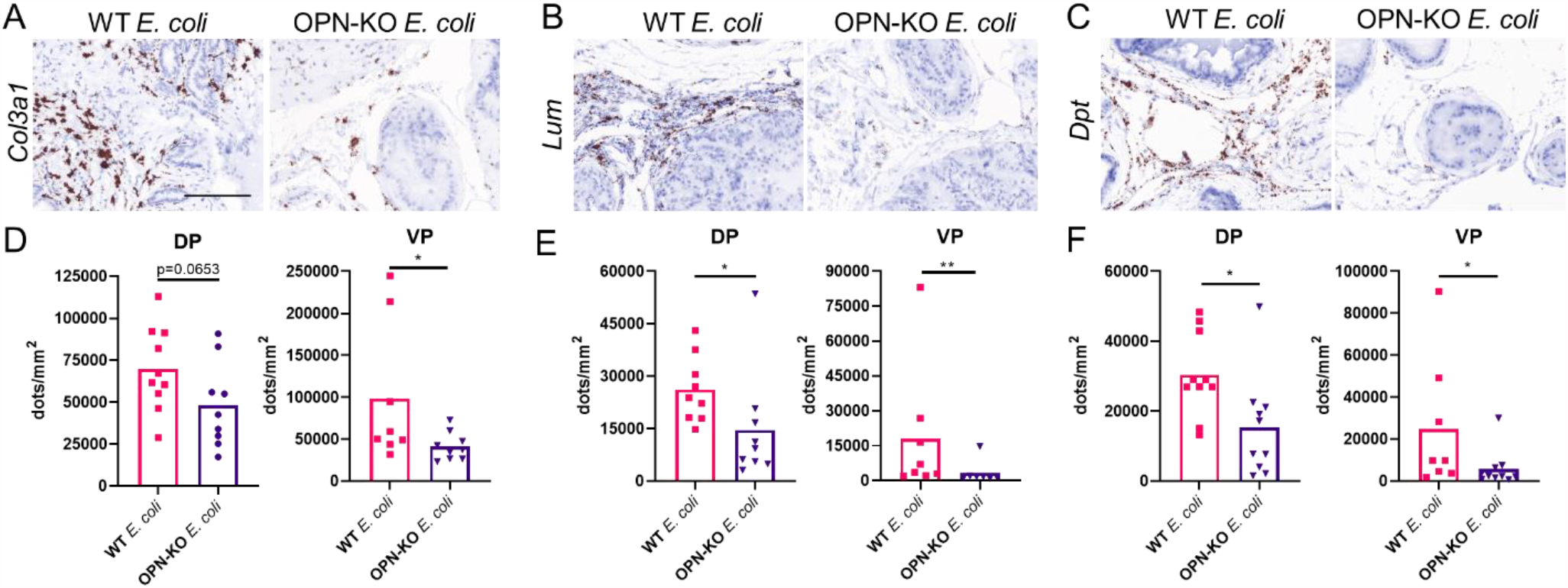
The pro-fibrotic genes, *Col3a1, Lum* and *Dpt*, are suppressed by OPN deficiency. Panels **A, B, C** are representative images and **D, E** and **F** show quantification of RNAscope staining of *Col3a1, Lum* and *Dpt*, respectively in the DP and the VP from *E. coli*-instilled WT and OPN-KO mice. Significance was tested using Mann-Whitney test. Images were captured at 40x magnification. Scale represents 100 µm. ^*^: p<0.05; ^**^: p<0.01.

## DISCUSSION

In recent years, substantial evidence has been accumulating on the crucial role of prostatic fibrosis in LUTS pathology, symptomology and therapeutic failure.^21,22,24^ However, since the etiology and molecular mechanism of fibrosis in the prostate is not yet clear, its diagnosis and treatment in LUTS patients are still out of reach. Prostatic fibrosis is triggered by inflammation, which occurs with high incidence in pathological prostate specimens.^12,42,43^ Bacterial and autoimmune experimental prostatic inflammation models both present with a fibrosis component^18,19,44^ which also supports the inflammatory origin of prostatic fibrosis. These findings inspired us to investigate how inflammatory signals initiate and exacerbate the fibrotic process in the prostate.

While characterizing a carrageenan-induced rat chronic inflammation model, we identified a robust increase in the expression of *Spp1* that encodes the pro-inflammatory and pro-fibrotic extracellular matrix protein and cytokine, OPN.^30^ Our further investigation revealed that OPN protein abundance is higher in surgically progressed human BPH tissue compared to incidental BPH^30^, potentially indicating a correlation of OPN levels with disease progression. Accordingly, we hypothesized that OPN translates inflammatory signals to fibrogenesis and thereby contributes to the pathogenesis of BPH/LUTS.

In the present study, we first characterized the tissue localization and expressional changes in SPP1/OPN in bacterial prostatic inflammation. While there was very limited expression of SPP1/OPN in saline-instilled controls, we found a robust induction in response to inflammation. This is consistent with previous reports showing that basal SPP1/OPN expression may be low but inducible by chemical^45^ or physical injury^28^ or by inflammation.^46^ OPN expression is upregulated in a variety of cells in response to stimuli, including immune cells^47^, fibroblasts^28^, epithelial cells^48^ or organ-specific cells such as alveolar epithelial cells^29^ or stellate cells.^49^ We have previously shown that OPN is expressed in prostate epithelial cells and fibroblasts, with epithelial cells producing higher basal OPN levels.^30^ M2 Macrophage-derived OPN has also been shown to stimulate epithelial proliferation.^33^ These results suggest that various cells can synthesize OPN in the prostate, although the majority of OPN is produced by inflammatory cells which are overrepresented in the *E. coli*-inflamed prostate.

To decipher whether systemic OPN expression is required for the development and/or resolution of prostatic inflammation and fibrosis, we instilled *E. coli* in WT and OPN-KO mice. The acute inflammatory grade and colonization rate were largely similar in WT and OPN-KO mice. However, most of the CD45+ immune cells and deposited collagen were cleared in the OPN-KO mice after 2 months. This suggest that OPN delays healing in the prostate after an inflammatory attack and potentially supports the transition from acute to chronic inflammation. Wound healing and the promotion of scar formation is an important role of OPN; when OPN expression is suppressed in skin injury models, collagen fibrils have reduced diameters and reduced immune cell infiltration to the site of injury leading to accelerated healing.^28^ Various studies explored the role OPN in inflammatory and fibrotic diseases and showed similar observations. OPN deficiency protected against aldosterone-induced inflammation, oxidative stress and interstitial fibrosis in the kidney.^50^ In thioacetamide-induced liver fibrosis, the rate of collagen deposition clearance was much higher in OPN-KO mice during a two-month healing phase.^27^ Mice deficient in OPN possessed less collagen-I and collagen-IV positivity in renal ischemia^51^ and less collagen, fibronectin, laminin and vitronectin levels in angiotensin-induced myocardial fibrosis.^52^

OPN may act on multiple levels to promote fibrosis. OPN increases collagen-I protein expression as shown in hepatocytes and in human lung fibroblasts.^29^ OPN increases α-smooth muscle actin levels providing collagen-producing myofibroblasts via integrinαvβ3 receptors and the phosphoinositide 3-kinase/phosphorylated Akt/nuclear factor kappa B (PI3K/pAkt/NFκB)– signaling pathway.^53^ Moreover, transgenic mice overexpressing OPN in hepatocytes develop spontaneous fibrosis^53^ highlighting that OPN is by itself sufficient to initiate the fibrotic process. In biliary epithelial cell (BEC)-hepatocyte co-culture, BECs release OPN that in turn increases the expression of the most profound pro-fibrotic factor, TGF-β1.^45^ OPN also stimulates fibroblast migration and proliferation by promoting the migration of fibroblasts to the site of injury to produce extracellular matrix^29^ and the activation of interstitial fibroblasts to myofibroblasts.^54^ Our previous studies using human prostate stromal and epithelial cell lines identified that OPN stimulates the stromal production of cytokines and chemokines, such as Il6, CXCl-1, -2, and -8 potentially exacerbating the inflammatory processes. It is possible that with the lack of OPN in the present study, acute inflammation is ameliorated and does not transform into chronic inflammation. Consequently, the activity and density of fibroblast and other potential collagen-producing cells (fibrocytes, myofibroblasts etc.) may decline and the tissue undergoes accelerated healing.

Despite OPN being expressed in most immune cells^55^, the volume of the initial inflammatory response and collagen deposition was not significantly impacted in OPN-KO mice. Our study, however, did not elucidate how OPN deficiency affects the distribution and activity of immune cell types, which may potentially contribute to the resolution of fibrosis OPN-KO seen two months after bacterial instillation. Macrophage migration towards cancer cells is reduced in OPN-KO mice^56^ and OPN also regulates macrophage polarization.^57^ In addition, OPN activates T helper 1 cells.^58^ A recent paper with IPF shows that *Spp1* is primarily expressed in myeloid cells in interstitial pulmonary fibrosis and IL-6 stimulates OPN expression in macrophages, which sensitizes and mobilizes fibroblasts toward other fibrogenic growth factors.^59^ Accordingly, it is likely that one major outcome of SPP1 deficiency is altered immune cell type distribution and activation, which possibility will be addressed in future studies.

To explore the transcriptional landscape of WT and OPN-KO prostates in chronic inflammatory conditions, we performed a bulk RNA-seq analysis on VPs. A robust expressional change was detected in WT mice instilled with *E. coli* that corresponds to the high degree of inflammatory and fibrotic processes in this group. Genes that were upregulated included collagens, collagen processing enzymes, non-collagenous ECM structural proteins, proteoglycans, cell surface glycoproteins, growth factors and their binding proteins and cytokines and chemokines. Most importantly, these results are the first to provide a comprehensive analysis of ECM gene expression signature that is responsible for the development of chronic fibrosis in the mouse prostate. Interestingly, high expression of key matrix proteins, such as *Col1a1, Col3a1, Fbn1* and *Dcn* and enzymes processing their assembly (*Lox, Loxl*s), co-occur with an increase in *Mmp*s (Mmp3, -7, -9, -12 and -23) that drive the clearance of the ECM. *Mmp*s degrade ECM molecules and therefore they are proposed to counteract increased ECM deposition and pro-fibrotic processes.^60^ In contrast, there is growing evidence that several *Mmp*s possess pro-fibrotic roles.^61^ In inflamed environments, *Mmp*s can modulate cytokines and chemokines and release. chemotactic fragments or activate growth factors to promote inflammation and fibrosis.^62^ Thus, their stimulation in our *E. coli*-induced inflammation model is not surprising and suggests an active tissue remodeling process.

From the 21 genes that were upregulated in WT *E. coli* and downregulated in OPN-KO *E. coli* in our RNA-seq dataset, we selected and confirmed similar expressional changes for four pro-fibrotic genes, *Lum, Col3a1, Dpt* and *Mmp3*, across all four experimental groups and two prostate lobes. Lumican is a proteoglycan important in the regulation of collagen fibril assembly^63^ but its expression is increased in fibrosis and *Lum* deficiency or suppression attenuates fibrosis.^64,65^ The increase in *Col3a1* expression and changes in collagen-I:collagen-III ratio alter collagen-I microstructure^66^ and are implicated to produce therapeutic resistance to corticosteroids in pulmonary fibrosis.^67^ Dermatopontin regulates fibronectin polymerization^68^ and enhances TGF-β activation.^69^ *Mmp3* is one of the many *Mmp*s that cleaves OPN to increase its adhesive and migratory activity.^70^ Mmp3 also promotes fibrosis by activating the β-catenin pathway and promoting epithelial-to-mesenchymal transition in idiopathic pulmonary fibrosis.^71^ According to our observation, the upregulation of these genes seems to represent both an increase in the number of cells expressing these genes, as well as an increase in expression. Suppression of these four genes in *E. coli*-instilled OPN-KO mice suggest that the deficiency in OPN contributes to a faster healing process where the deposition of ECM proteins is suppressed.

Our RNA-seq dataset also identified *Cyp2b10* as a gene upregulated by OPN deficiency. In the liver, the enzyme encoded by its human homolog, *CYB2B6*, is involved in xenobiotic metabolism but also in the hydroxylation and deactivation of testosterone and androstendione.^72^ *CYB2B6* was shown to be expressed in prostate epithelial cells and suppressed in prostate cancer cells, its downregulation correlates with poor prognosis and its overexpression inhibits proliferation in LNCaP prostate cancer cells.^73^ Inflammatory cytokines, especially IL-6, generally downregulate the expression of cytochrome P450 enzymes^74^, but how their suppression contributes to inflammation and fibrosis in the prostate is unclear and needs further investigation.

In summary, we showed that OPN is a pro-inflammatory and pro-fibrotic factor that delays the resolution of fibrosis in bacteria-instilled mouse prostates. The deficiency of OPN leads to the substantial reduction of ECM-producing cells consistent with the downregulation of various matrix-related genes. As OPN-targeting therapies are currently under development for various cancers and fibrotic diseases ^75,76^, our study identifies that they may be beneficial in the treatment of LUTS patients, especially when symptoms are associated with significant prostatic fibrosis.

## METHODS

### Mice

Animal experiments were conducted under the protocols approved by the University of Wisconsin (UW) Animal Care and Use Committee (ID: M005570-R01-A04). Breeding pairs of Spp1^tm1Blh^/J and male C57BL/6J mice for wild type (WT) controls were obtained from The Jackson Laboratory (Bar Harbor, ME). The Spp1^tm1Blh^/J strain was originally donated by Liaw et al.^35^ and were inbred at the UW Biomedical Research Model Services for no more than 3 generations. This strain was backcrossed to C57BL/6J for more than 10 generations by the donating lab and the live colony was established in C57BL/6J mice using cryopreserved sperm at The Jackson Laboratory. Animals were maintained on a strict 12:12-h light-dark cycle in a temperature- and humidity-controlled facility.

### Bacterial culture and transurethral instillation

The UTI89 uropathogenic *Escherichia coli* strain used for transurethral instillation was isolated from a patient with cystisis and transformed with a PCOMGFP plasmid providing kanamycin resistance and green fluorescent protein expression.^18,77,78^ One colony of UTI89 *E. coli* was used to inoculate 100 ml Luria-Bertani (LB) broth and incubated by shaking at 200 rpm at 37°C for 16 hours. Cells were harvested by centrifugation at 1400g for 15 min. and resuspended in sterile PBS. Optical density (OD) was adjusted to 0.8. UTI89 or PBS was instilled via a transurethral catheter (OD 0.8, 100 µl) two times, 3 days apart, to 8-week-old WT or OPN-KO mice. Necropsy was performed and tissues were collected one or 8 weeks after the first instillation. Free catch urine was collected 24 hours after the second instillation and colony forming units (CFU) were determined as described earlier.^18^

### Void spot assay

Void spot assays (VSAs) were used to determine changes in urinary function as described previously.^79^ Mice were placed individually in cages securely lined with 15.9 × 27.1-cm filter paper for 4 hours (from 10 AM to 2 PM) with food but without water. The filter papers were air dried and imaged with an Autochemi AC1 Darkroom ultraviolet imaging cabinet (UVP, Upland, CA) as described earlier.^18^ Images were imported into ImageJ and void spots were quantified with VoidWhizzard.^80^ Only urine spots in the 0-0.1 cm^2^ category were analyzed. Overlapping spots were analyzed based on the software’s algorithm. Mice were acclimatized to the VSA conditions by performing the assay once without analyzing the data and then obtaining the basal VSA parameters from the subsequent experiment.

### Picrosirius red staining and polarized imaging

Picrosirius red staining was performed and analyzed as previously described.^23^ Briefly, tissues were deparaffinized, hydrated and stained with 0.1% Sirius red F3BA in saturated aqueous picric acid (Sigma-Aldrich, St. Louis, MO) for 1 hour. Nuclei were counterstained with Weigert’s hematoxylin (Sigma-Aldrich), dehydrated, and mounted. Birefringence was captured under circular polarized light and the density of 4 different colors (green, yellow, orange, red, indicative of elevating collagen bundle thickness) was quantified. The ratio of individual colors was expressed relative to the number of total positive pixels. Total collagen was calculated as the proportion of positive pixels relative to the tissue area calculated in ImageJ with thresholding of a bright field image captured at the same position as with the polarized light.

### Hematoxylin-Eosin (H&E) staining and Immunohistochemistry (IHC)

H&E staining was performed by a standard method using Shandon Instant Hematoxylin and Eosin-Phloxine (Thermo Fisher Scientific, Waltham, MA) multichrome stain. For IHC, sections were de-paraffinized and hydrated. Antigen retrieval was acquired in a decloaking chamber (Biocare Medical) using citrate buffer pH 6.0. Endogenous peroxidases and non-specific binding sites were blocked with Bloxall (Vector Labs) and horse serum (2.5% or 10%) or Rodent Block, respectively. Primary antibodies anti-COL1A1 (NBP30054, Novus Biologicals, Centennial, CO, Lot: SH1120a 1:500 dilution, Lot: SH1219R 1:300 dilution), anti-CD45 (ab10558, Abcam, 1:2000 dilution) and anti-OPN (AF808, R&D Systems, Minneapolis, MN, 1:100 dilution) were added overnight. An HRP-conjugated horse Anti-Rabbit IgG Polymer (MP-7401, Vector Laboratories) or rabbit anti-goat IgG antibody (1:250 dilution, Bethyl Laboratories, Montgomery, TX) was used as secondary for 30 minutes and the signal was developed using the SignalStain DAB Substrate Kit (Cell Signaling, Danvers, MA).

### RNAscope

RNAscope (*in situ* hybridization) was performed using probe sets specific to mouse *SPP1* (435191, target region: 2 – 1079 bp,), Col3a1 (455771, target region: 873 – 1711), Lum (480361, target region: 254 – 1336), Dpt (561511, target region: 2 – 1045) and Mmp3 (480961, target region: 739 – 1798) with the RNAscope® 2.5 HD Assay-Red or –Brown detection system (Advanced Cell Diagnostics, Newark, CA). Reaction specificity was tested with a positive control probe and a negative control with only a probe diluent instead of the probe.

### Image analysis

IHC tissues were imaged with a Mantra 2 Quantitative Pathology Workstation (Akoya Biosciences) with a 20x objective. Three representative images were taken per tissue. Area not containing tissue was removed (CD45+ and OPN) or only the stroma was selected (for COL1A1 density) with inForm software using trainable tissue segmentation. Optical density (OD) or number of positive cells was calculated by the software. CD45-positive cells were counted by hand in the chronic inflammation group. Cell counts were normalized to tissue area.

RNAscope staining was captured with a 40x objective (6 images/tissue) and staining area was determined using the Inform software thresholding function. Individual particle size (averaged 0.8 micron) was used to calculate particle number from total positive area and normalized to tissue area.

### RNA isolation, library preparation, RNA-seq and analysis (not final, waiting for feedback from Biotech center)

Ventral lobes were thawed and the RNA was stabilized in RNAlater (Thermo Fisher Scientific) for 1 hour. Total RNA was isolated from ventral lobes using the Qiagen RNeasy Micro kit with on-column DNAse digestion (Germantown, MD). Library preparation, quality check, RNA sequencing and bioinformatics analysis were performed at the UW-Madison Biotechnology Center. RNA quantity was measured with NanoDrop and assayed on an Agilent 2100 Bioanalyzer. The cDNA library was prepared by the TruSeq Stranded Total RNA (Gold) Library Prep with after polyA enrichment and rRNA reduction. Libraries were quantified using Qubit DNA HS kit and assayed on Agilent Tapestation DNA 1000 system. Libraries were sequenced on a NovaSeq 6000 platform as paired-end, 150bp reads and fastq files were generated with bcl2fastq2 conversion software v2.20 using reads meeting Q30 in >80% of reads. The trimming software Skewer^81^ (v0.1.123) was used to trim Illumina adaptors from the fastq files. Trimmed reads were aligned to the mouse genome GRCm38.p5 (accession NCBI:GCA_000001635.7) using the splice-junction aware read aligner STAR (v2.5.3a). Based on ensembl version 88 annotation (March 2017), mapped paired-end reads for genes were counted in each sample using RSEM v1.3.0 (RNASeq by Expectation Maximization), described by Li and Dewey^82^. Samples were normalized by the method of trimmed mean of M-values (TMM), where the product of this factor and the library sizes defines the effective library size^83^. Independent filtering was performed prior to differential expression analysis, requiring a threshold of least 1 transcript count per million in at least 3 samples, ignoring any prior group assignment. Analysis of differentially expressed genes was performed using EdgeR package^84^ (v3.16.5). Statistical significance of the negative-binomial regression test was adjusted with a Benjamini-Hochberg FDR correction at the 5% level^85^. Volcano plots were generated using VolcaNoseR.^86^

### Statistical analysis

All statistical calculations were performed in Graphpad Prism (Graphpad Software, San Diego, CA). One-way ANOVA was used if the Brown-Forsythe and Bartlett’s tests were significant followed by Tukey’s post hoc analysis. Otherwise, the non-parametric Kruskal-Wallis test with Dunn’s multiple comparison test for groups more than 2 was performed. To compare two groups, we used two-tailed t-test when the F-test was significant. Otherwise the statistical difference was determined by Mann-Whitney test. Figure 2A was created using Biorender.com.

## Availability of materials and data

The RNA-seq data and analysis discussed in this publication have been deposited in NCBI’s Gene Expression Omnibus^87^ and are accessible through GEO Series accession number GSE179655 (https://www.ncbi.nlm.nih.gov/geo/query/acc.cgi?acc=GSE179655). The datasets generated during and/or analyzed during the current study are available from the corresponding author on reasonable request.

## Acknowledgements

The authors thank the University of Wisconsin Biotechnology Center Gene Expression Center, DNA Sequencing Facility and Bioinformatics Resource Center for providing library preparation, RNA sequencing and analysis services. We thank Emily Ricke, Christian Ortiz Hernandez and Livianna Myklebust and Drs. Teresa Liu, Jordan Vellky and Dalton McLean for experimental assistance and advice. This study was supported by grants from the National Institutes of Health including K12 DK100022-06 and K01 DK127150-01 (to P.P.), U54 DK104310 and R01 ES001332 (to W.A.R. and C.M.V.).

## Author contributions

P.P., W.A.R. and C.M.V. conceived of and designed the experiments and wrote the paper. P.P., A.J., K.O.S., E. S., K.S.U. performed and analyzed the experiments and generated the figures.

H.R. and M.C. provided training to perform the experiments and conceptual support. All authors reviewed the manuscript.

