## Supplemental Information for "Osteopontin deficiency leads to the resolution of prostatic fibrosis and inflammation"

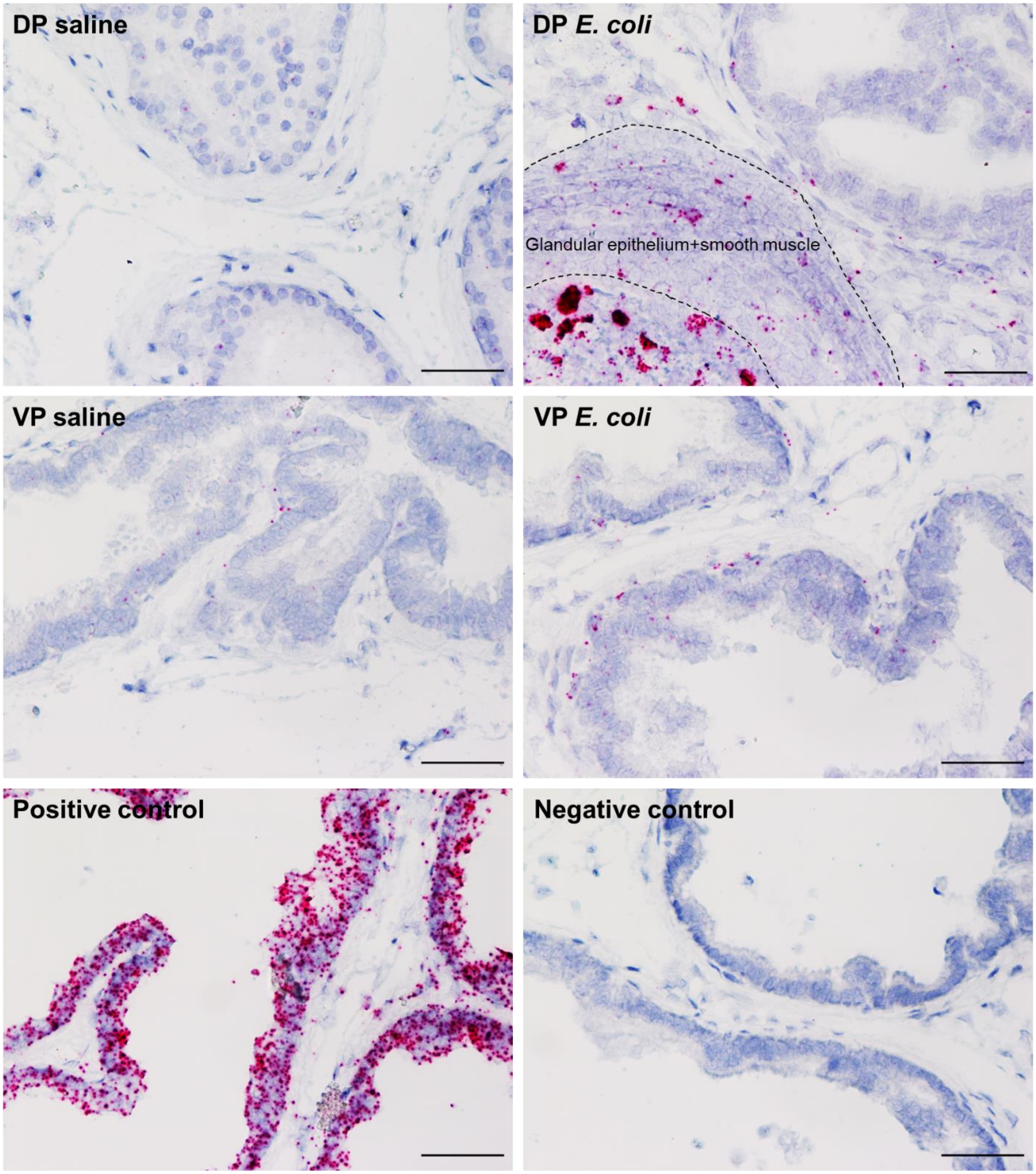


**Supplemental Fig. 1** SPP1 expression is upregulated in the DP in E. coli-instilled mice. SPP1 expression appears to be upregulated in multilayered prostate ducts as well as in the lumens that are occupied by immune cells. The VP only contained sporadic upregulation of SPP1 expression. The positive control probe was provided by Advanced Cell Diagnostics and targeted a gene that has medium expression level across all cells. For the negative control, no probe was added. Scale represents 100 µm.

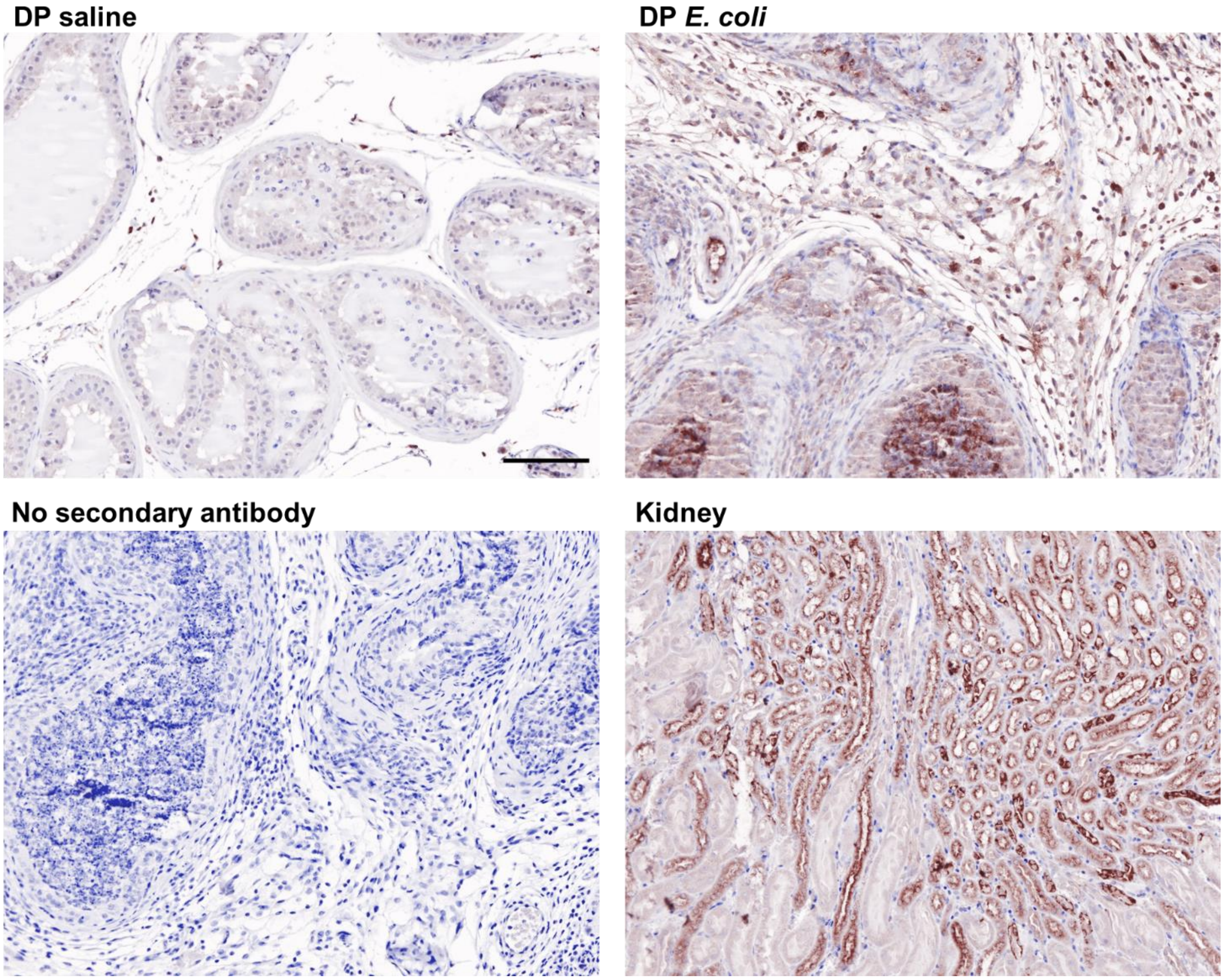


**Supplemental Fig. 2** OPN protein expression is upregulated in the DP in E. coli-instilled mice. OPN expression appears to be upregulated in multilayered prostate ducts as well as in the lumens that are occupied by immune cells. Images were captured at 20x magnidication. Scale represents 100 µm.

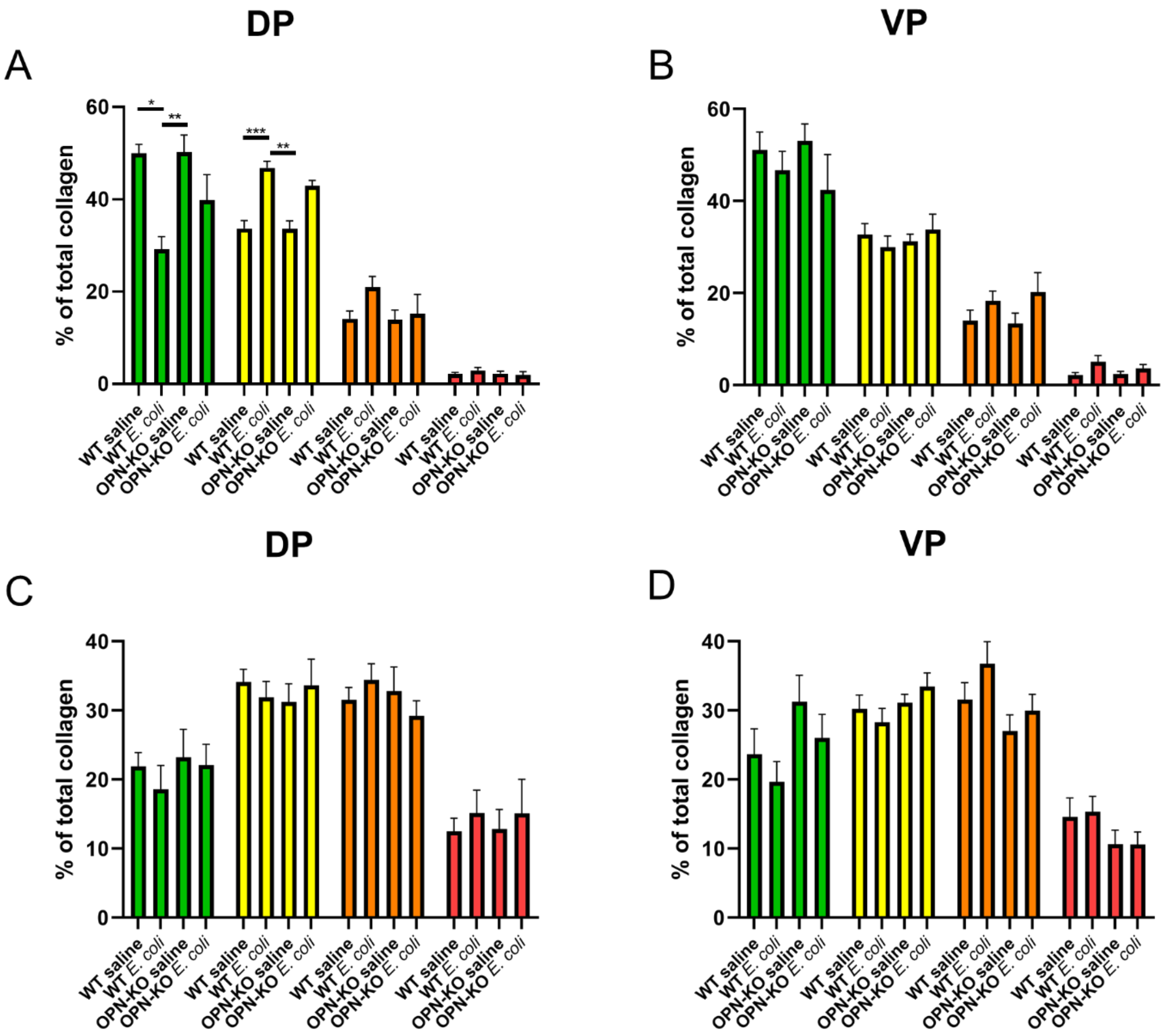


**Supplemental Fig. 3** The distribution of collagen polarization colors change similarly in WT and OPN-KO *E. coli*-instilled dorsal prostates. One week after the bacterial instillation (A and B), we found significant decrease in green color and increase in the proportion of yellow fibers in the DP, but not in the VP, in WT mice which indicates an increase in collagen thickness. There was no significant alteration in the representation of polarization colors after two months (C and D). Tissues were stained with PSR and images were taken with a circularly polarized filter. Significance was determined by the Kruskal-Wallis non-parametric test. *: p<0.05; **: p<0.01; ***: p<0.001.

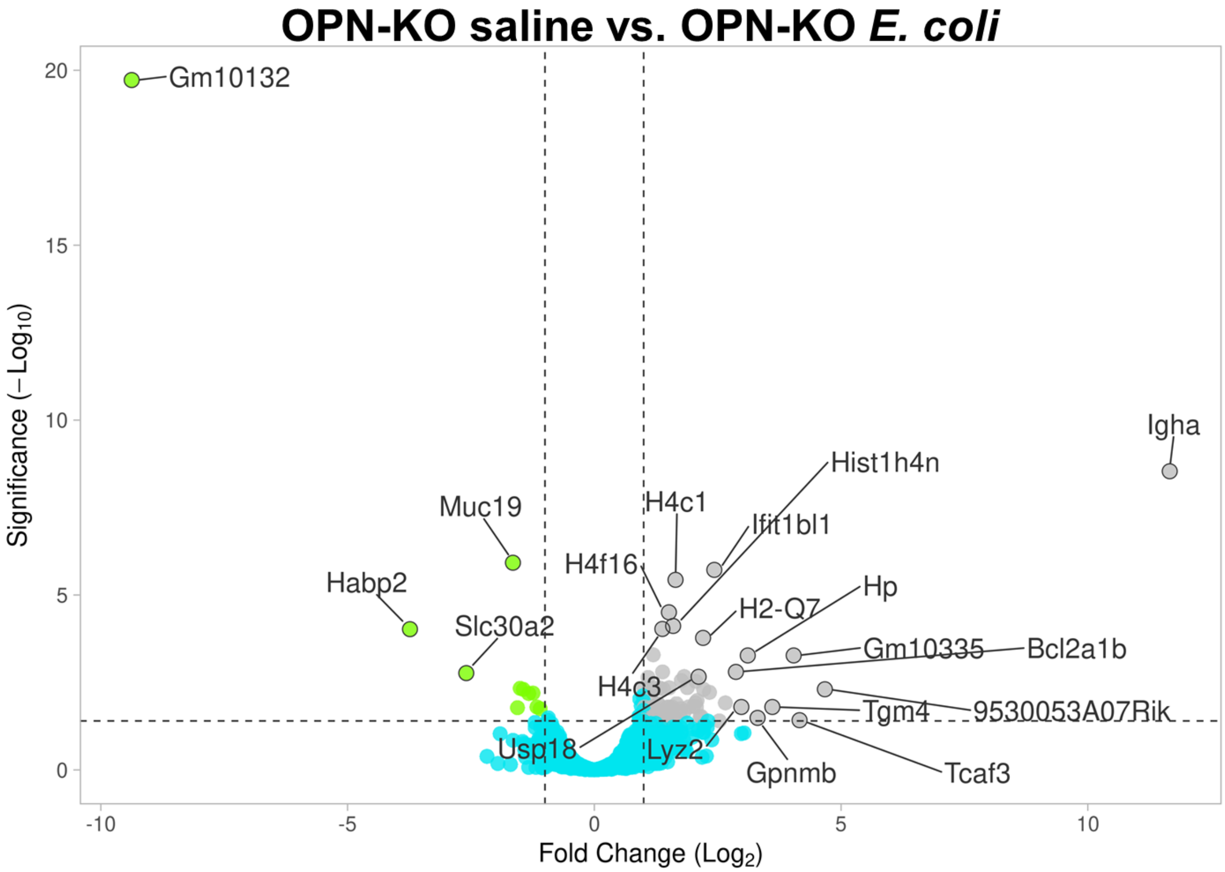


**Supplemental Fig. 4**. OPN deficiency prevents *E. coli*-induced expressional changes in genes associated with inflammation and fibrosis. The 20 top hits determined by Manhattan distance with the VolcaNoseR application are labelled.

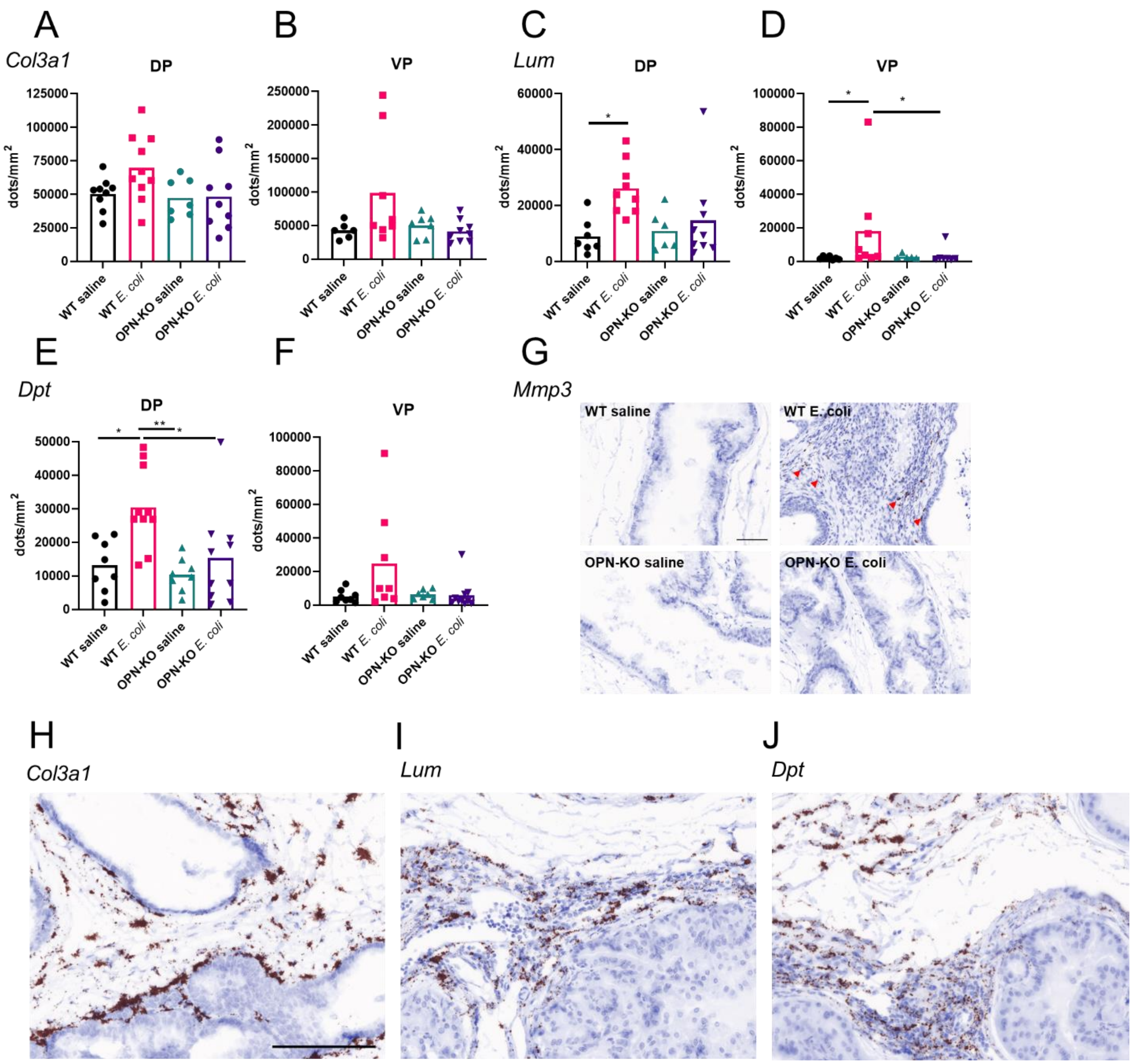


**Supplemental Fig. 5**. RNAscope analysis of pro-fibrotic genes, *Col3a1* (Panels **A** and **B**), *Lum* (Panels **C** and **D**), *Dpt* (Panels **E** and **F**) and *Mmp3* (Panel **G**) in the DP and the VP across all experimental groups. Quantification of mRNA abundance for *Col3a1*, *Lum* and *Dpt* was performed using the Mantra 2 Quantitative Pathology Workstation. Due to the limited expression of *Mmp3* in uninflamed tissues, representative images are shown instead of quantification. Representative images for *Col3a1*, *Lum* and *Dpt* are shown in Panels **H**, **I** and **J**, respectively. Significance was determined by the Kruskal-Wallis non-parametric test. Images were captured at 40x magnification. Scale represents 100 µm. *: $p<0.05$; **: $p<0.01$.
